# Visceral Fat Inflammation and Fat Embolism are associated with Lung’s Lipidic Hyaline Membranes in COVID-19 patients

**DOI:** 10.1101/2021.10.30.466586

**Authors:** Georgia Colleluori, Laura Graciotti, Mauro Pesaresi, Angelica Di Vincenzo, Jessica Perugini, Eleonora Di Mercurio, Sara Caucci, Patrizia Bagnarelli, Cristina M. Zingaretti, Enzo Nisoli, Stefano Menzo, Adriano Tagliabracci, Annie Ladoux, Christian Dani, Antonio Giordano, Saverio Cinti

**Affiliations:** Center for the Study of Obesity, Department of Experimental and Clinical Medicine, Marche Polytechnic University, via Tronto 10A, Ancona, Italy; Section of Experimental Pathology, Department of Clinical and Molecular Sciences, Marche Polytechnic University, via Tronto 10A, Ancona, Italy; Section of Legal Medicine, Department of Bioscience and Public Health, Marche Polytechnic University, via Tronto 10A, Ancona, Italy; Section of Microbiology, Department of Bioscience and Public Health, Marche Polytechnic University, via Tronto 10A, Ancona, Italy; Center for Study and Research on Obesity, Department of Medical Biotechnology and Translational Medicine, University of Milan, via Vanvitelli 32, Milan, Italy; Université Côte d’Azur, CNRS, Inserm, iBV, Faculté de Médecine, 06107 Nice Cedex 2, France

**Keywords:** SARS-CoV2, visceral adipose tissue, obesity, inflammation, pulmonary embolism, pneumonia

## Abstract

**Background:** Visceral obesity is a critical determinant of severe coronavirus disease-2019 (COVID-19). Methods: In this study, we performed a comprehensive histomorphologic analysis of autoptic visceral adipose tissues (VAT), lungs and livers of 19 COVID-19 and 23 non-COVID-19 subjects.

**Results:** Although there were no between-groups differences in body-mass-index and adipocytes size, higher prevalence of CD68+ macrophages in COVID-19 subjects’ VAT was detected (p=0.005) and accompanied by crown-like structures presence, signs of adipocytes stress and death. Consistently, human adipocytes were successfully infected by SARS-CoV2 *in vitro* and displayed lower cell viability. Being VAT inflammation associated with lipids spill-over from dead adipocytes, we studied lipids distribution employing Oil-Red-O staining (ORO). Lipids were observed within lungs and livers interstitial spaces, macrophages, endothelial cells, and vessels’ lumen, features suggestive of fat embolism syndrome, more prevalent among COVID-19 individuals (p<0.001). Notably, signs of fat embolism were more prevalent among obese (p=0.03) independently of COVID-19 diagnosis, suggesting that such condition may be an obesity complication, exacerbated by SARS-CoV2 infection. Importantly, all infected subjects’ lungs presented lipids-rich (ORO+) hyaline membranes, formations associated with COVID-19-related pneumonia, present only in one control with non-COVID-19 pneumonia.

**Conclusions:** This study describes for the first time novel COVID-19-related features possibly underlying the unfavorable prognosis in obese SARS-CoV2-infected-subjects.

## Introduction

Since December 2019, the severe acute respiratory syndrome coronavirus 2 (SARS-CoV2), responsible for the development of coronavirus disease 2019 (COVID-19), has spread globally resulting in a worldwide health crisis that caused over four million deaths (1). The lung is a crucial target organ not only due to the severe bilateral pneumonia observed in 15-30% of hospitalized patients (2, 3), but also because it is the site from which the infection spreads to blood vessels, heart, gut, brain, and kidneys (4). Published data support interstitial fibrosis with alveolar hyaline membrane (HM) formation as the main underlying histopathologic event responsible for pneumonia and acute respiratory syndrome distress (5, 6). The reasons for HM bilateral expression, histogenesis, and sudden clinical appearance during COVID-19 early stages are not completely understood (7).

The severity of COVID-19 is strictly associated with the presence of comorbidities (8); while obesity alone is responsible for 20% of COVID-19 hospitalizations, obesity in combination with type 2 diabetes and hypertension accounts for 58% (9). Obesity and impaired metabolic health are in fact strongly associated with COVID-19 unfavorable prognosis and pose also young patients at higher risks (10, 11). Importantly, visceral obesity increases the risk of COVID-19-related complications, independently of age, gender, body mass index (BMI), total and subcutaneous adipose tissue areas (12-15). Visceral obesity is in fact strongly associated with chronic low-grade inflammation, blood hypercoagulability, impaired metabolic health, and higher risk of cardiovascular events, all risk factors for COVID-19 severity (8, 11, 15-17). Visceral adipose tissue (VAT) excessive expansion is paralleled by adipocytes hypertrophy, death, and lipids spill-over, phenomena resulting in macrophages infiltration, crown-like structures (CLS) development and inflammation, in turn contributing to the obesity-related complications (18-20). The elevated adipocytes ACE2 expression in obesity (21), receptor exploited by SARS-CoV2 for cell entry, has been often speculated as a possible pathophysiological mechanism responsible for obesity-related COVID-19 severity (8, 22, 23). However, although obesity has been strongly associated with COVID-19 severity (but not higher infection rates), original articles comprehensively analyzing adipose tissue samples belonging toCOVID-19 subjects and providing direct evidence of SARS-CoV2 infection are lacking (15). In our preliminary study, we observed the presence of fat embolism in a COVID-19 subjects with obesity, a phenomenon that we hypothesized could derive from adipose tissue stress induced by SARS-CoV2 and explain COVID-19 severity in obesity (22). In the present study we perform for the first time a comprehensive histomorphological assessment of visceral adipose tissue, lung, and liver autoptic samples belonging to COVID-19 and non-COVID-19 subjects, and specifically focusing on tissues lipids distribution. We observed novel SARS-CoV2-related histopathological features *i*.*e*., visceral adipose tissue inflammation, signs of fat embolism and lung’s hyaline membranes of lipidic nature, possibly contributing to the severity of COVID-19 among subjects with visceral obesity.

## Materials and methods

### Study Approval

We followed the report “Research ethics during COVID-19 pandemic: observational, and in particular, epidemiological studies” published by the Italian *Istituto Superiore di Sanità* on May 2020 (Rapporto ISS COVID-19, n. 47/2020) (37). Given the observational (cross-sectional, case-control) nature of our study which was conducted on autoptic specimens and did not entail neither an intervention, nor the collection of subject’s sensitive information, we have not obtained an informed consent. Our study did not entail any physical risk for the subjects. In Italy, the evaluation of non-pharmacological observational studies is not governed by the same normative references provided for the evaluation of clinical trials and observational studies concerning drugs. Furthermore, as reported in the above report (37) in the section dedicated to our type of study *in conditions of pandemic and therefore of high risk for the communities, some administrative steps may be abolished*. Therefore, our Institutional Review Board does not require an ethical approval for studies conducted on autoptic specimens and not collecting personal or sensitive data.

### Study subjects and tissue sampling

Autoptic lung, liver, and visceral adipose tissue samples of 49 subjects were collected at the Department of Legal Medicine of the Ospedali Riuniti of Ancona between March 2020 and May 2021. Twenty-four subjects were affected by COVID-19, while the remaining 25 were not and died for different reasons. SARS-CoV2 infection was assessed in all subjects by RT-PCR tests on nasopharyngeal swab. Subjects were included in the analyses only if their lung’s samples were well preserved such that a high-quality histological assessment could be performed. We hence analysed 19 COVID-19 cases and 23 controls. Among the studied subjects, 15 had documented respiratory conditions -*i*.*e*., pneumonia, dyspnoea, respiratory distress-(10 COVID-19 and 5 controls), 15 had documented hypertension (7 COVID-19 and 8 controls), 11 suffered from type 2 diabetes (6 COVID-19 and 5 controls) and 10 from cardiovascular diseases (2 COVID-19 and 8 controls). Visceral adipose tissue was sampled from the omentum and mesentery region. Lungs were extensively sampled across central and peripheral regions of each lobe bilaterally. A median of seven tissue blocks (range five to nine) were taken from each lung. Liver samples were collected from the right and left lobe.

Samples were sliced into different pieces to be studied by light microscopy (LM) and transmission electron microscopy (TEM). A comprehensive methodological description for such methodologies has been described elsewhere (38).

### Immunohistochemistry and morphometric analyses

The collected visceral (omental) adipose tissue, lung and liver autopsies were fixed overnight at 4°C in 4% paraformaldehyde. Samples were then embedded in paraffin to be studied by LM and to perform immunohistochemistry and morphometric analyses. For each sample, 3 µm paraffin sections were obtained and used for immunohistochemical analyses. A comprehensive description of the protocol has been described elsewhere (38). To detect the presence of CD68+ macrophages in VAT samples, we used CD68 (Dako #M0814; dilution 1:200; antigen retrieval method by citrate buffer pH6) antibody. To study SARS-CoV2 presence in VAT, we used the SARS-CoV2 nucleocapsid (Invitrogen #MA-17404) and spike protein (Sino Biological #40150-T62) antibodies at different dilutions. The same antibodies were used to detect the virus on infected VeroE6 at dilution: 1:1000 for nucleocapsid protein and 1:100 for the spike protein. To assess antibody specificity, negative control in which primary antibody was omitted were always included in each set of reaction. Tissue sections were observed with a Nikon Eclipse E800 light microscope. For morphometric purposes, for each paraffin section, 10 digital images were acquired at 20X magnification with a Nikon DXM 1220 camera. CD68 positive macrophages widespread in VAT parenchyma and those organized to form CLS were counted in all images. For each subject the number of total macrophages and the density of CLS/104 adipocytes were counted with the ImageJ morphometric program (RRID:SCR_003070). Adipocytes’ area was measured in all patients by counting 100 adipocytes for each paraffin tissue section using ImageJ.

### Histochemical staining

For Oil Red-O (ORO) staining samples were cryoprotected in 30% sucrose overnight, embedded in the optimal cutting temperature (OCT) compound medium, and then sliced to obtain 7 µm-thick cryosections by Leica CM1900 cryostat (Vienna, Austria). ORO staining was then performed on lungs (43) and liver (n=9) cryosections. In brief, dried cryosections were first placed in 60% isopropanol, then in filtrated Oil-Red O working solution (15 minutes at room temperature) and briefly washed again in 60% isopropanol and lastly in H_2_O. Tissue slices were then counterstained with hematoxylin and cover with a coverslip using Vectashield mounting medium (Vector Laboratories). Lung and liver tissues organization and morphology were also studied by hematoxylin & eosin (H&E) staining on paraffin sections. Lung’s hyaline membranes presence and characterization were performed on paraffin sections by H&E, periodic acid-Schiff and Masson trichome staining.

### Transmission electron microscopy

For ultrastructural analyses, 3-mm thick VAT (n=4), lung (n=7) and liver (n=1) samples were further fixed in 2% glutaraldehyde-2% paraformaldehyde in 0.1 M phosphate buffer (pH 7.4) and post-fixed in Osmiun Tetroxide 1% then embedded in epoxy resin for TEM studies as described elsewhere (38). Cell pellets from the *in vitro* studies were similarly fixed in 2% glutaraldehyde-2% paraformaldehyde in 0.1 M phosphate buffer (pH 7.4) for one hour at room temperature and then embedded in epoxy-resin. A MT-X ultratome (RMC; Tucson) was used to obtained ultrathin sections (∼70 nm). Ultrastructural characterization was performed on all samples using a CM10 Philips transmission electron microscope (Philips, Eindhoven, The Netherlands, http://www.usa.philips.com).

### Statistical analysis

Between-group comparisons for linear and categorical variables were determined by unpaired two-tailed Student’s t-test and Chi-square test, respectively. Group differences were considered significant when p<0.05. Data in graph are expressed as mean ± SEM. Statistical analyses were performed with Prism 6.0 (GraphPad Software Inc., La Jolla, CA) and IBM SPSS Statistics Data Editor (v.24).

### SARS-CoV2 infection in VeroE6

Vero E6 cells were cultured in Dulbecco’s modified Eagle medium (DMEM, Euroclone, Milano, Italy), supplemented with 10% fetal calf serum (FCS Euroclone) and antibiotics/antimycotic (100 U/ml penicillin, 100 µg/ml streptomycin, 0.25 µg/ml amphotericin B) at 37°C, 5% CO_2_ in a humidified atmosphere (90%), as described previously (39). Cells were maintained in 75 cm^2^ tissue culture flasks. The day before infection, a confluent monolayer was trypsinized, and 1,5 × 10^6^ cells were seeded in every 8 flasks (25 cm^2^). Confluent monolayers were infected with SARS CoV-2 (78952 isolate, accession no. MT483867) (40) at a multiplicity of infection (MOI) of 3.29·10^5^. After 2 hours of incubation, the medium containing the inoculum was removed, the cells were washed twice, and fresh medium was added, which was collected after 6, 12, 24 and 48 h for viral genome quantification and replaced with 2 ml of fresh culture medium to allow scraping of the infected monolayer. Uninfected cell monolayer controls were treated as infected ones. Cell suspensions (2ml) were subsequently centrifuged at 800 rpm for 5 minutes. Aliquots of infected supernatants, collected as above, were analyzed using RT-qPCR assay as described elsewhere (40). Briefly, 5 µl of RNA extracted from 140 µl of infected supernatants were run together with a calibration curve, obtained from 10-fold dilutions of a standard plasmid certified and quantified by a supplier (2019-nCoV Positive Control, nCoVPC, 85 IDT) and negative control, applying a protocol described by CDC (https://www.fda.gov/media/134922/download).

### In vitro studies on hMADS

Ethical Approval: Human adipocytes progenitors -Aps-(hMADS cells) were isolated from adipose tissue, as surgical scraps from surgical specimen of various surgeries of young donors, with the informed consent of the parents. All methods were approved and performed in accordance with the guidelines and regulations of the Centre Hospitalier Universitaire de Nice Review Board.

#### Cell Differentiation

hMADS cells were maintained and differentiated as previously described (41). They will be further referred to as hMADS-adipocytes. They were routinely tested for the absence of mycoplasma. Treatments and biological assays were carried out in duplicates on control or differentiated hMADS cells from day 4 to 18.

#### Gene expression analysis

Total RNA was extracted using the TRI-Reagent kit (Euromedex, Soufflweyersheim, France) and reverse transcription (RT) was performed using MMLV reverse transcriptase (Promega, Charbonnieres, France), as recommended by the manufacturers. All primer sequences are described in the supplementary section. Real-time PCR assays were run on an ABI Prism One step real-time PCR machine (Applied Biosystems, Courtaboeuf, France). Normalization was performed using *36B4* as a reference gene. Quantification was performed using the comparative Ct method. The results are shown as mean ± standard error of the mean (SEM), with the number of experiments indicated. Statistical significance was determined by *t-*tests BiostaTGV (INSERM and Sorbonne University, PARIS, France). Probability values <0.05 were considered statistically significant and are marked with a single asterisk, <0.01 with double asterisks and <0.001 with triple asterisks. Sequences for the primers used in this study *ACE2* (FW 5’-AGAACCCTGGACCCTAGCAT -3’; REV 5’-AGTCGGTACTCCATCCCACA -3’); *BASIGIN* (FW: 5’-CAGAGTGAAGGCCGTGAAGT -3’; REV: 5’ACTCTGACTTGCAGACCAGC-3’); *NRP1 (FW:* 5’-GGGGCTCTCACAAGACCTTC 3’; REV: 5’-GATCCTGAATGGGTCCCGTC -3’); *CSTL* (FW: 5’-CTGGTGGTTGGCTACGGATT -3’; REV: 5’-CTCCGGTCTTTGGCCATCTT -3’); FURIN (FW:5’-CTACAGCAGTGGCAACCAGA-3’; REV:5’-TGTGAGACTCCGTGCACTTC-3’); *36B4 (FW:* 5’-CTACAACCCTGAAGAAGTGCTTG -3’; REV: 5’-CAATCTGCAGACAGACACTGG -3’); *DPP4* (SINO biologicals Inc. #HP100-649 (Eschborn, Germany)

#### hMADS Sars-CoV2 infection

hMADS and hMADS adipocytes cells were infected with viral stock of SARS-CoV2 (EPI_ISL_417491), at a 50% Tissue Culture Infectious Dose (TCID_50_) of 2000 TCID_50_/ml for 2 hours at a temperature of 37°C. Following incubation, the medium containing the inoculum was removed, the cells were washed twice, and the medium was supplemented with different specific compounds. Supernatants were collected at 24, 48, 72, 96 hours for viral genome quantification and medium renewal was performed at each sampling time. Uninfected cell monolayer controls were treated as the infected ones. Supernatants, collected as above, and cell pellets, collected at 96 hours post-infection, were analyzed using RT-qPCR as described in the VeroE6 cell section.

#### *Cell Viability Assay* (MTT Assay)

The effect of SARS-CoV2 infection on cell viability of hMADS adipocytes was measured using the metabolic dye [4,5-dimethylthiazol-2-yl]-2,5-diphenyl tetrazolium (MTT) (Sigma, St. Louis, MO, USA). Briefly, hMADS cells were seeded in 96 well plates at a density of 4,500 cells/cm^2^, differentiated and then infected with viral stock of SARS-CoV2 for 2h at 37 °C. Following the incubation with the virus, cells were placed in supplemented medium. Time-course analyses of cell survival were determined at 24, 48, 72 and 96h. After the incubation period, the media were replaced with 100 µL MTT (0.5 mg/mL) dissolved in PBS and incubated for 3 h. MTT-containing medium was removed and 100 μl of dimethyl sulfoxide (DMSO) was added to dissolve formazan crystals formed by live cells. Absorbance was subsequently measured at 570 nm using a BioTek Synergy HTX microplate reader (BioTek, Winooski, VT, USA). Results were expressed as percentages of viable cells relative to uninfected controls.

#### Nuclear morphology analyses

Alterations in nuclear morphology were determined by assessment of nuclear staining using fluorescent stains and fluorescent microscopy (42).

For these experiments, hMADS adipocytes were differentiated in 2-well Lab-Tek Chamber Slides (Nalge Nunc International, Naperville, IL, USA), washed with PBS pH 7.4 and fixed with 10% paraformaldehyde in PBS for 10 min at RT. After washing with PBS, nuclear staining was performed with Hoechst. Finally, cells were airdried and cover-slipped using Vectashield mounting medium (Vector Laboratories, Burlingame, CA, USA) and analyzed by fluorescent microscopy. The number of altered nuclei were counted (in the field displaying nuclear fragmentation, nuclear condensation) and divided by the total number of nuclei and multiply by 100. Observations were carried out by Lucia IMAGE 4.82, Laboratory Investigations Morphometric Analyses.

*Lipid droplet size* (µm^2^) was measured on SARS-CoV2 infected hMADS adipocytes and in untreated controls. For this purpose, we used a drawing tablet and a morphometric program (Nikon LUCIA IMAGE, Laboratory Imaging, version 4.61; Praha, Czech Republic). hMADS adipocytes were examined with a Nikon Eclipse Ti-S inverted light microscope (Nikon Instruments S.p.A, Calenzano, Italy), and digital images were captured at 20X with a Nikon DS-L2 camera (Nikon Instruments S.p.A, Calenzano, Italy). Five random fields were analyzed and at least 1700 lipid droplets were measured for each sample, and the difference between infected and non-infected cells was assessed by unpaired t-test. Similarly, the quantitative assessment of the material extruded from the hMADs was calculated using the same microscope and software and expressed as the number of vacuoles extruded from the cells on the total cell amount.

## Results

Autoptic VAT, lung and liver samples belonging to 49 subjects were collected and screened to be included in the study. Forty-two subjects were considered suitable for the study (good-preservation for histomorphologcal analyses), 19 of which died due to COVID-19-related bilateral pneumonia (COVID-19 group), while the remaining 23 died for different reasons (control group). Subjects’ characteristics, including gender, age, BMI, comorbidities, and cause of death are reported in supplementary table 1 and 2. SARS-CoV2 infection was assessed by RT-qPCR performed on nasal pharyngeal or pharyngeal swab samples. Study population mean age was 65.0±14.3 years old, BMI was 29.0±5.4 kg/m^2^ with 35.7% of patients suffering from obesity (BMI≥30.0 kg/m^2^), and 45.2% being overweight (BMI≥25.0 kg/m^2^). Thirty-five % of the population was composed of woman (n=15). There were no significant differences in mean age (COVID-19: 69.5±11.0 vs controls: 61.0±16.0 years old; p=0.09) and BMI (COVID-19: 30.0±5.0 vs controls: 28.1±5.6 kg/m^2^; p=0.62) between our study groups.

Unequivocal signs of chronic, low-grade inflammation in both COVID-19 and control subjects with a BMI≥25.0 kg/m^2^ were observed in VAT samples (Fig.1A). However, although there were no between-groups differences in BMI and VAT adipocytes size (Fig.1B), higher prevalence of CD68+ macrophages (Fig.1C) and a trend for higher presence of CLS (Fig.1D) were evidenced in COVID-19 patients compared to controls, suggesting higher SARS-CoV2-induced VAT inflammation. Other inflammatory cells were represented mainly by lymphocytes, but their number was negligible in all investigated cases.

We then assessed whether the higher VAT inflammation in COVID-19 patients was associated with adipocytes death. Perilipin 1 (PLIN1) immunohistochemistry is a reliable method for identification and quantification of dead adipocytes (18, 24). However, in the present study, all samples display PLIN1 negative adipocytes, probably due to the autoptic nature of specimens. We hence performed a morphologic and ultrastructural study to assess VAT adipocytes stress and death. Electron microscopy showed signs of adipocytes death in proximity of CLS in both COVID-19 and controls subjects with a BMI≥25 kg/m^2^, a finding consistent with previous studies documenting obesity-related adipocytes death (25). However, COVID-19 subjects VAT was rich in stressed and dead adipocytes (Fig. 1E-F) also in areas lacking CLS and seemingly normal at light microscopy. In line with the observed widespread death, cell remnants were evidenced in closed proximity of dying adipocytes, while free lipid droplets were often found in fat interstitial spaces (Fig.1F and 1G). Notably, large lipid vacuoles were also observed: *i*. inside endothelial cells belonging to capillaries adjacent to free lipid droplets (Fig.1H and Fig.1I); *ii*. extruding from endothelial cells into the capillary lumen (Fig.1I); *iii*. in the lumen of VAT capillaries (Fig.1J); *iv*. in macrophages near interstitial free lipid droplets (data not shown). In addition, several clusters of lipid-rich structures were found into the lumen of venules belonging to mesenteric fat samples (Fig.1K). In summary, the in-depth ultrastructural analyses of VAT autoptic samples belonging to COVID-19 subjects revealed the widespread presence of free lipid droplets (likely deriving from dead adipocytes) inside the capillary lumen, all features underlining a condition able to generate fat embolism syndrome (FES) (26).

**Figure 1.**
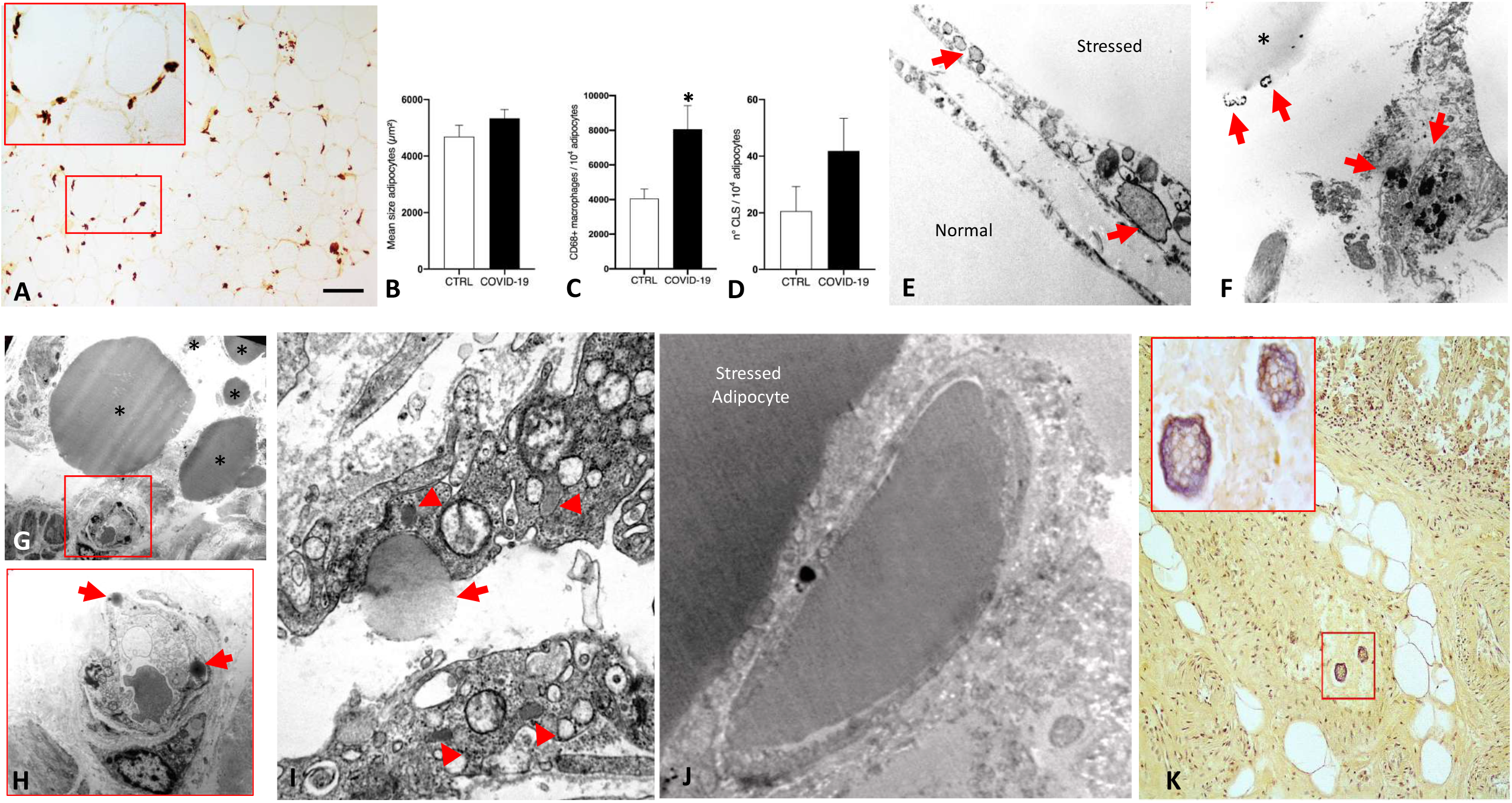
Visceral adipose tissue inflammation and fat embolism in COVID-19 subjects. (**A)** Light microscopy (LM): representative immunohistochemistry of visceral adipose tissue infiltrated by CD68+ macrophages (in brown); inset shows an enlargement of the squared area. **(B)** Visceral adipose tissue adipocytes area, **(C)** number of CD68+ macrophages per 10^4^ adipocytes, and **(D)** number of crown-like structures (CLS) per 10^4^ adipocytes in COVID-19 vs control subjects. Asterisk (*) indicates p<0.05. **(E)** Transmission electron microscopy (TEM): normal adipocyte adjacent to a stressed adipocyte showing dilated endoplasmic reticulum (arrows). **(F)** TEM: dead adipocytes and interstitial free lipid droplets (*); arrows indicate adipocytes remnants. **(G)** TEM: free lipid droplets of variable size were frequently found in COVID-19 subjects (asterisks). **(H)** Enlargement of squared area in G showing lipid droplets inside endothelial cells (arrows). **(I)** TEM: enlargement of a capillary from a COVID-19 subject showing a lipid droplet extruding into the capillary lumen (arrow), note the abundant Weibel-Palade bodies denoting increased blood hypercoagulability (arrowheads). **(J)** TEM: a capillary filled with embolic fat near a stressed adipocyte. **(K)** LM: mesenteric fat sample showing lipid-rich embolic material in a vein (squared area, enlarged in inset). Morphometric data are expressed as means±SE. Scale Bar: A=100 µm, E=0,8 µm, F=2,5 µm, G=10 µm, H=3 µm, I=1,5 µm, J=0,8 µm, K= 35 µm.

We then aimed at assessing whether the observed VAT alterations were associated with SARS-CoV2 local-tissue presence or if they were a consequence of the systemic infection. Although SARS-CoV2 ability to infect human adipose tissue has been frequently speculated (8, 12, 17, 22), direct evidence of such phenomenon has not been documented in the literature (15), with only one study reporting the presence of the virus in mediastinal fat (27). While SARS-CoV2 genomic RNA, nucleocapsid and spike proteins were not detectable in VAT samples of COVID-19 subjects, virus-like structures with morphology and size resembling the those present in SARS-CoV2 infected VeroE6 cells (Fig. 2A) were found in the cytoplasm of stressed adipocytes (Fig.2B). Furthermore, the presence of ribosome-like clusters, described in virus-infected cells (28) was evidenced in both, visceral adipocytes belonging to COVID-19 subjects (Fig. 2C and 2D) and SARS-CoV2-infected VeroE6 (Fig. 2E). In addition, confronting cisternae, ribosome lamella complex and annulate lamellae, typical of several pathologic conditions including virus infection (29), were observed in VAT adipocytes belonging to COVID-19 subjects (Suppl. Fig.1A-D) and in SARS-CoV2 infected VeroE6 (Suppl. Fig.1E), but not in uninfected controls. Next, to provide direct evidence of SARS-CoV2 ability to infect human adipocytes, leading to cell stress and death, we infected differentiated human multipotent adipocytes (hMADS) (Fig. 2F-H) and studied SARS-CoV2 kinetics *in vitro*. The growth kinetics of SARS-CoV2 was determined as viral load (copies/ml) in the supernatants collected after 24-, 48-, 72- and 96-hours post-infection (Fig. 2F). While SARS-CoV2 genomic RNA was detectable in both, differentiated and undifferentiated hMADS at the first timepoints post-infection (24 and 48 h), it could be detected only in mature adipocytes at later timepoints (72 and 96 h) (Fig. 2F). Consistently, SARS-CoV2 genomic RNA was also detected in the hMADS adipocytes pellet after 96-hours of infection (Fig. 2G). Importantly, infected hMADS adipocytes displayed lower cell viability (Fig. 2H), higher prevalence of pyknotic nuclei (Fig. 2I-K) and smaller lipid droplet size - suggestive of cell delipidation and stress-compared to uninfected controls (Fig. 2L). Furthermore, in line with these data, evidence of increased material extrusion from infected cells were evidenced by light microscopy (p<0.05) and strongly suggested massive cell delipidation induced by SARS-CoV2 (Suppl.1F-H). We hence performed a time-course analyses of hMADS expression of putative SARS-CoV2 receptors (Fig. 2L) and proteases (Fig. 2M) in presence or absence of the adipogenic differentiation cocktail (at 4, 7, 14 and 18 days). *ACE2* receptor was expressed at very low levels in both differentiated and undifferentiated hMADS, even though we used specifically designed primers holding a 100.92% efficiency. On the other side, *BASIGIN* receptor was preferentially detected in differentiated hMADS which displayed an increased expression after 14 days. The receptor *NEUROPILIN* 1 was expressed by undifferentiated cells. Concerning proteases expression, while differentiated hMADS expressed the protease *FURIN*, the undifferentiated ones preferentially expressed *DPPIV*. The expression of *CATHEPSIN L* did not differ between the two conditions, while we did not detect *TMPRSS2* in both differentiated and undifferentiated hMADS (data not shown).

**Figure 2.**
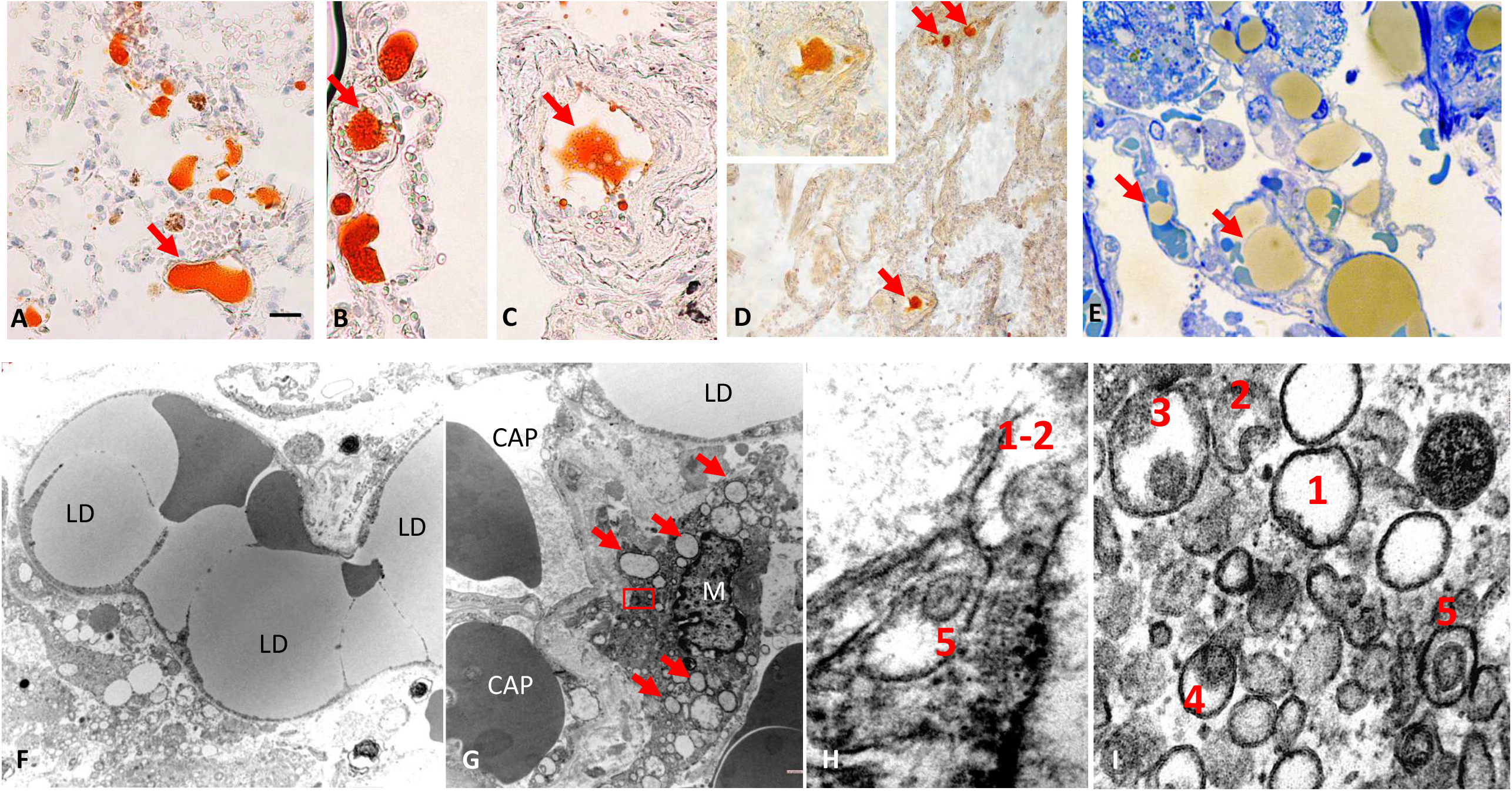
SARS-CoV2 in visceral adipose tissue and hMADS. (**A)** Transmission electron microscopy (TEM): Vero E6 infected cell showing several virions into the rough endoplasmic reticulum (RER), some indicated by arrows. Inset: enlargement of squared area. (**B)** TEM: adipocyte from the visceral adipose tissue (VAT) depot of a COVID-19 subject showing several virions into RER (arrows). Inset: enlargement of squared area. **(C)** TEM: VAT of a COVID-19 subject showing an adipocyte (Ad) with two large ribosome-like clusters (dotted lines) in the cytoplasm. **(D)** Enlargement of squared area in C showing ribosome-like cluster and a virion-like structure into the dilated RER (arrow). **(E)** TEM: SARS-CoV2 infected VeroE6 cells showing a ribosome-like cluster (squared area), enlarged in the inset. **(F)** SARS-CoV2 infection kinetic in undifferentiated and differentiated hMADS. SARS-CoV2 genomic RNA detected in the supernatant at different timepoints, expressed as copies (cps)/ml. **(G)** SARS-CoV2 quantification in supernatant and cell pellets of hMADS infected cells. **(H**) MTT viability assay in SARS-CoV2 infected and uninfected hMADS adipocytes shows lower cell viability in the first compared to the last at 24- and 96-hours post-infection (p<0.05). **(I)** Percentage of pyknotic nuclei in hMADS adipocytes at 96h post-infection compared to uninfected controls (p<0.05). **(J)** Hoechst nuclear staining showing pyknotic nuclei (arrows) in differentiated hMADS adipocytes. **(K)** Lipid droplets average area (µm^2^) in differentiated hMADS 96h post infection compared to uninfected controls (p<0.0001). Expression of putative SARS-CoV2 receptors **(L)** or proteases **(M)** assessed by RT-qPCR and normalized for the expression of *36B4* mRNA. Expressions were measured in cells that received (red bars) or did not receive (blue bars) the differentiation cocktail for the indicated number of days. The means ± SEM were calculated from three independent experiments (*ACE2, BSG, NRP1, CSTL*) or four independent experiments (*FURIN, DPP4*), with determinations performed in duplicate (*p<0.05, ** p<0.01). Scale Bar: A, B =200 nm, C=500 nm D=100 nm E=180 nm, F=120 µm, G=70 µm, H=5 µm.

Given our preliminary data (22) and the widespread lipid droplets presence in the capillary lumen of VAT, also evidenced in some mesenteric adipose depots, we then studied lipid distribution in lung samples employing Oil-Red O staining (ORO: lipid-specific histochemistry). Lipids were evidenced within lungs alveolar septa, interstitial spaces, endothelial cells and vessel’s lumen and in alveolar and interstitial macrophages (Fig. 3A-D), all features confirmed by light and electron microscopy (Fig. 3E-F) and suggestive of fat embolism (21).

**Figure 3.**
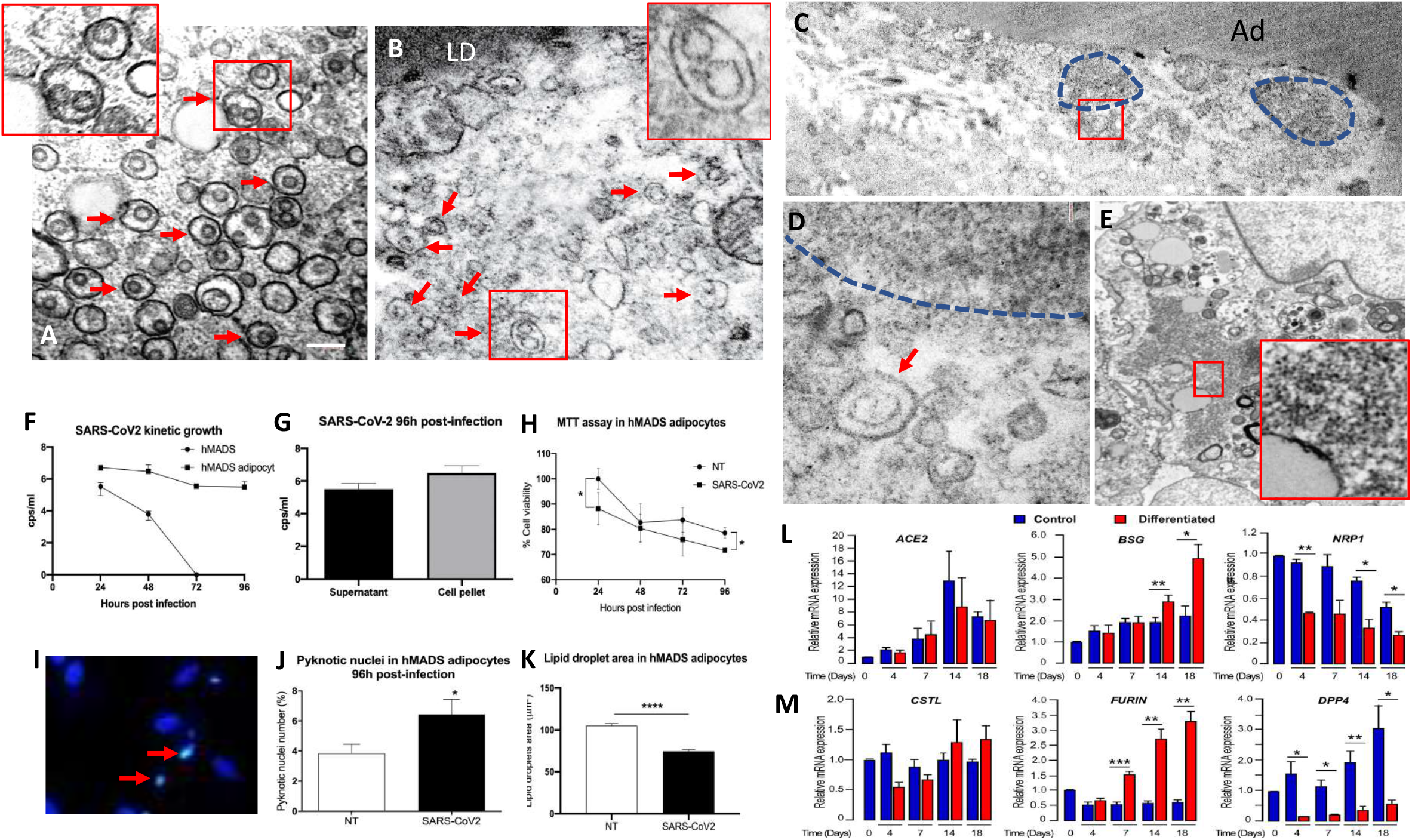
Embolic lipid droplets and SARS-CoV2 virions in lung of COVID-19 subjects. **(7)** Light microscopy (LM): representative histochemistry for fat (Oil-Red O) showing the lipid nature of vacuoles (orange-red) in the vascular lumen (arrows) and lung septa of different COVID-19 subjects. **(E)** LM: resin embedded, toluidine-blue stained tissue. Large free lipid droplets (yellow) are evident into the capillaries lumen in alveolar septa (arrows). **(F)** Transmission electron microscopy (TEM): showing lipid droplet (LD) into an alveolar septum mixed with erythrocytes. **(G)** TEM: alveolar macrophage (M) in a COVID-19 subject. Note: diffuse dilated rough endoplasmic reticulum (RER) denoting cellular stress (arrows) **(H)** TEM: enlargement of the squared area in G showing two virions at stages 1-2 and 5 of the reproductive cycle into the dilated RER similar to what observed in **(I)** TEM: (1 to 5) stages of reproductive cycle of SARS-CoV2 virions in VeroE6 infected cells. Reference in the main text. Scale Bar: A, B, C=20 µm, D=140 µm E=8 µm, F=1,5 µm, G=1 µm, H=70 nm I=65 nm.

Lung’s fat embolism was not exclusive of, but more prevalent among COVID-19 subjects as compared to controls (100% vs 53%; p<0.001). Signs of fat embolism were in fact more prevalent among individuals with obesity than in those with a BMI≤30 kg/m^2^ (93% vs 63%, p=0.03), independently of COVID-19 diagnosis. Consistently, all subjects with type 2 diabetes (T2DM) had fat embolism. Of note, electron microscopy observation revealed several structures with size and morphology compatible with those of SARS-CoV2 viruses (6) in pneumocytes, endothelial cells and macrophages, the last of which displayed disseminated, dilated endoplasmic reticulum denoting cellular stress (25, 30) and signs of virus presence only in COVID-19 subjects (Fig. 3G-H). Furthermore, we also evidenced also two virions at early and late stages of reproductive cycle (31) into the dilated endoplasmic reticulum (Fig. 3H) comparable with those revealed in infected VeroE6 in Fig. 3I. Importantly, septal capillaries very often contained large amounts of fibrin, with some of them lining by fibrin-thrombotic material only in COVID-19 individuals’ lungs (data not shown). Several Weibel-Palade bodies, signs of activated coagulative phenomena (29), were observed also in capillary endothelial cells belonging to COVID-19 subjects (data not shown).

Unexpectedly, the ORO technique evidenced also positively stained alveolar structures reminiscent of hyaline membranes (Fig. 4A). The presence of hyaline membranes was then confirmed by hematoxylin and eosin, by Mallory and periodic acid-Schiff staining (data not shown). All COVID-19 subjects presented lung’s hyaline membranes, which were on the other side detected only in one control subject (BMI 21.3 kg/m^2^) who died of pneumonia (p<0.0001). Interestingly, this last subject displayed fainted lung’s hyaline membrane positivity for ORO staining, suggesting a lower lipidic composition. This finding is consistent with other reports describing hyaline membrane presence in pneumonia (7). Importantly, ORO positive lipid droplets and lipid-rich macrophages were often enclosed into the hyaline membranes lining the alveolar surface (Fig. 4B-D). Several aspects suggesting a direct role of embolic fat in hyaline membranes formation were observed. Specifically, free lipid droplets occupying the alveolar space and lining and spreading on the alveolar surface were observed (Fig. 4E-H). The presence of lung’s hyaline membranes of lipidic nature was associated with visceral adipose tissue inflammation (8.0±5.4 vs 3.7±1.8 CD68+ macrophages/10 adipocytes in subjects with and without hyaline membranes, respectively) and exclusive of COVID-19 cases (Suppl. Fig. 2).

**Figure 4.**
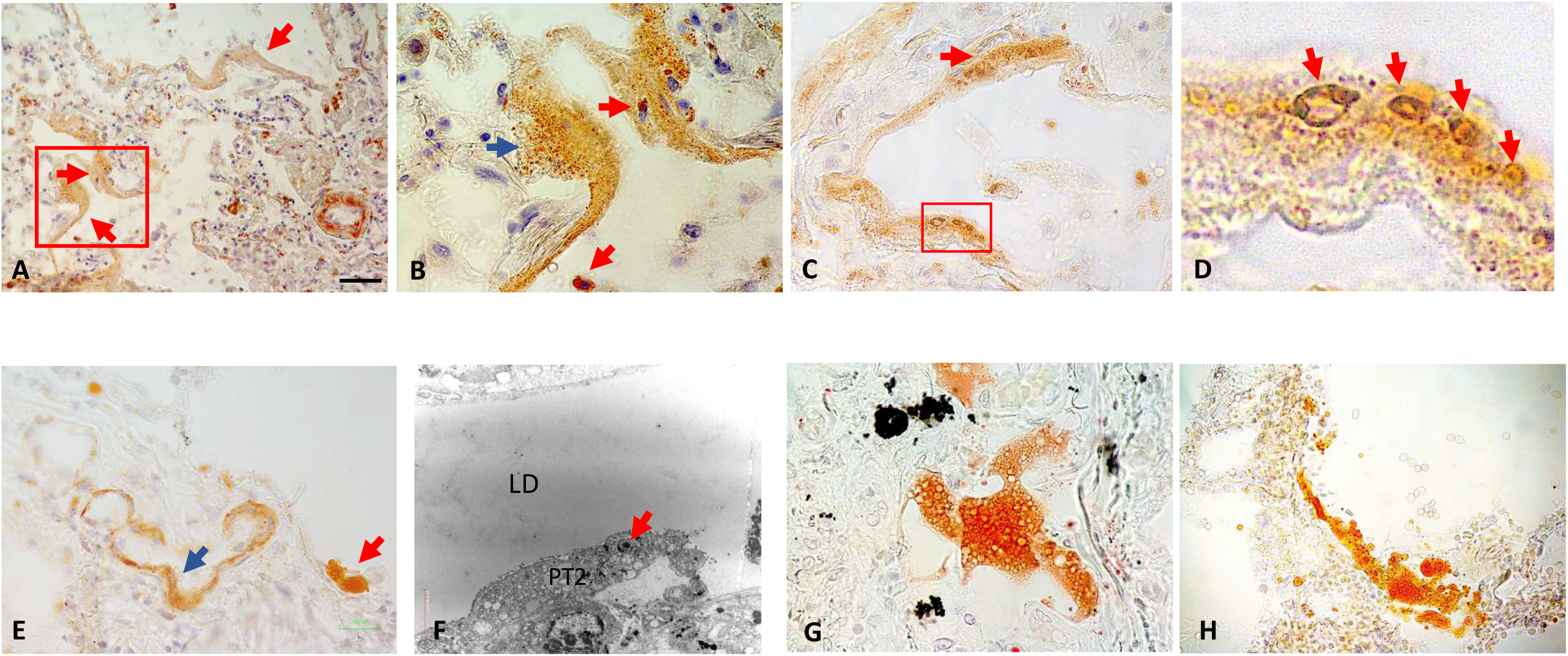
Oil-Red O-stained lung of COVID-19 subjects showing hyaline membranes morphology and composition. **(A)** Light microscopy (LM): hyaline membranes lining alveolar surfaces (arrows) at low magnification. **(B)** LM: enlargement of squared area in A showing the microvacuolar nature of ORO+ hyaline membrane (blue arrow). Lipid rich macrophages free in the alveolar space (red arrows) and inside hyaline membranes (blue arrows) **(C)** LM: vacuolar aspect of ORO+ hyaline membranes’ lipids (arrow and squared area). **(D)** LM: enlargement of squared area in C. Arrows indicate lipid vacuoles. **(E)** LM: ORO+ large, free lipid vacuole lining the alveolar surface (red arrow) near a hyaline membrane (blue arrow). **(F)** TEM: free lipid droplet lining the alveolar surface composed by pneumocytes type II (PT2) with classic surfactant granules (arrow). **(G)** LM: ORO+ lipid vacuole spreading on the alveolar surface (possible early stage of lipid diffusion). **(H)** LM: ORO+ lipid vacuoles possibly contributing to hyaline membranes development (later stage). Scale Bar: A and E= 50 µm, B=7 µm, C=10 µm, D=2 µm, F=3 µm, G=25 µm, H=35 µm.

Lastly, since the embolic material from abdominal visceral tissues should necessarily pass through the liver parenchyma to reach the lung, we exploited the ORO staining technique to study liver samples belonging to 9 COVID-19 and 8 control subjects. Liver autoptic samples showed focal, macrovescicular steatosis with lipid droplets of very variable size (Suppl. Fig. 3A), consistent with other studies conducted on COVID-19 subjects (32). In particular, signs consistent with fat embolism, i.e., presence of free lipid droplets into hepatic sinusoids (Suppl. Fig. 3B) and into the vessels’ lumen (Suppl. Fig. 3C-D), as well as clusters of lipid-rich structures in the portal vein (Suppl. Fig. 3D) were observed in COVID-19 subjects, a finding that confirmed the embolic nature of hepatic fat droplets, and that support what observed in VAT samples. In summary, 8/9 COVID-19 subjects with documented lung fat embolism displayed signs of hepatic fat embolism as well. On the other side, we observed hepatic embolism in an elevated percentage of control subjects (6/8), possibly due to the elevated prevalence of visceral obesity among these investigated cases.

## Discussion

This is the first study investigating the ultrastructural features of VAT among COVID-19 subjects and assessing lipid distribution in lungs and liver samples by histomorphology. Our data support the presence of higher local VAT inflammation and higher prevalence of fat embolism and lipidic hyaline membranes formations in the lungs of subjects dead due to COVID-19 compared to control individuals’ dead for different reasons. In addition, our data support SARS-CoV2 ability to infect human adipocytes *in vitro*. Considering the strong association between COVID-19 related complications and obesity, especially with visceral adipose content excess (10-15), the comprehension of the biological phenomenon at the basis of such association holds critical clinical implication in the era of the COVID-19 pandemic.

Our study provides the first evidence of higher local VAT inflammation among COVID-19 subjects, independently of obesity status and support COVID-19-induced exacerbation of obesity-related inflammation, a novel finding consistent with studies reporting higher systemic inflammation among infected patients (17). Adipocyte’s inflammation is associated with adipocytes stress, death and lipids release in the extracellular space (18, 19, 24, 25). We hence studied adipocytes features by TEM and revealed the presence of the typical signs of cellular stress, together with clear features of lipids’ spill-over from suffering adipocytes. Lipids were in fact detected in the extracellular space, inside endothelial cells, inside the capillary lumen, and extruding from endothelial cells into the capillary lumen, all features indicative of fat embolism.

Although virus like structures were evidenced by TEM in the same VAT depots, the lack of SARS-CoV2 detection by qPCR did not allow us to conclude that such inflammation, cellular stress and death were all related to the presence of this virus. It is in fact possible that the described VAT features were secondary to the systemic inflammation induced by COVID-19 or due to the presence of different viruses within the depot. On the other side, we were able to demonstrate that SARS-CoV2 can infect human adipocytes even though neither adipocytes, nor adipocytes progenitors gathered all the known molecular requirements for the virus entry (expression of all known virus proteases and receptors). This set of data is in part consistent with other findings and suggest that additional, not yet characterized, receptors and proteases may be exploited for this purpose (15, 33).

Considering the widespread lipid droplets presence in the capillary lumen of VAT and considering our preliminary data (22), we studied lipid distribution in lung and liver samples and confirmed the presence of fat embolism. Interestingly, we noticed similar lipid-like structures also in lung’s images from other reports on COVID-19 subjects, reason for which we believe it is worth performing further in-depth analyses on available samples (5, 6, 34).

Fat embolism was prevalent among, but not exclusive of, subjects with COVID-19; it was in fact detected also among subjects with obesity independently of SARS-CoV2 infection. These data are not surprising given that adipocyte’s death and release of lipids are both phenomena occurring in obesity (18, 24, 25). This finding provides the first evidence pointing out fat embolism as a complication of obesity (and obesity plus T2DM), determined by adipocytes death and possibly exacerbated by the COVID-19-induced inflammatory status. Importantly, studying lung’s lipid distribution, we unexpectedly revealed the presence of lipidic hyaline membranes, formation strongly contributing/associated to COVID-19 related interstitial fibrosis and pneumonia (6). Hyaline membranes were present in all COVID-19 subjects and in only one control who died for pneumonia, a finding consistent with other reports describing hyaline membrane presence in pneumonia (7). Our histomorpholgic assessment revealed several aspects indicative of a direct role of embolic fat in hyaline membranes formation. Consistently, the presence of lung’s hyaline membranes of lipidic nature was associated with visceral adipose tissue inflammation but was exclusive of COVID-19 cases.

In summary, in our case series, although fat embolism may be present in condition of obesity and T2DM independently of COVID-19, the embolic-derived pulmonary lipidic material contribute to the formation of hyaline membranes only in the case of COVID-19 related pneumonia, a novel finding that holds critical clinical implications and deserves further investigation. Furthermore, these data provide significant insight into hyaline membrane nature, as their formation process has not been characterized yet (35). Additional studies investigating the hyaline membranes nature of non-COVID-19-related pneumonia are required to detail such histopathological feature.

Collectively our data reveal higher local VAT inflammation in COVID-19 subjects and SARS-CoV2 ability to infect human adipocytes, both elements widely speculated but never demonstrated in the literature (15, 23). In addition, we provide the first evidence that supports fat embolism as a complication of obesity, likely determined by adipocytes death and exacerbated by the COVID-19-induced inflammatory status. Lastly, we reveal for the first time the presence of lung’s lipidic hyaline membranes among all infected subjects, a novel COVID-19-related histopathological feature associated with visceral adipose tissue inflammation and fat embolism. Consistently, fat embolism displays similar signs and symptoms as the ones observed in COVID-19, in line with a recently published case report (36). Differential diagnosis, when fat embolism and COVID-19 are suspected, is hence critical for proper patients’ care. Based on our findings, the assessment of fat embolism symptoms is mandatory in the context of the COVID-19 pandemic, especially among patients with pulmonary symptoms, obesity and high waist circumference, signs of elevated visceral adipose accumulation. Such complex clinical status should be therefore adequately assessed and properly addressed. Our data hold critical clinical implication in the context of obesity disease and the COVID-19 pandemic and need to be confirmed by additional studies with a larger sample size.

## Supporting information

Supplemental Figue 2

Supplemental Figue 3

## Acknowledgment

This study was funded by Fondo Integrativo Speciale per la Ricerca from the Italian Ministry of University and Research; grant number: FISR2020IP_05217, and supported by Progetti di Rilevante Interesse Nazionale (PRIN 2017, #2017L8Z2) and by Cariplo Foundation to E.N. (grant 2016-1006).

## Author Contribution

GC, LG, MP, AG, and SC: study conceptualization. GC, LG and SC: study coordination. MP and AT: collected autoptic samples and clinical data. GC, MP and ADV histological studies on autoptic samples and cell cultures. CMZ, LG and SC: electron microscopy studies. JP, EDM, AL and CD: *in vitro* studies on hMADS. SC, PB, SM: SARS-CoV2 infection for the *in vitro* studies. GC, LG, MP, JP, EN, SM, AG and SC: data analyses and interpretation. All authors approved the final version of the manuscript and take responsibility for its content.

---

**Suppl. Fig.1: Representative transmission electron microscope images of visceral adipose tissue from COVID-19 subject**

**(A)** Rough endoplasmic reticulum (RER) confronting cisternae in endothelial cell (arrowhead in A, enlarged in inset). **(B)** RER confronting cisternae in endothelial cell comparable to those found in infected VeroE6 cells (compare with E). **(C)** Ribosome-lamella complex in a capillary (arrow, enlarged in upper inset). **(D)** Annulate lamellae found in a lung’s macrophage. **(E)** RER confronting cisternae in SARS-CoV2 -infected Vero-E6 cell. **(F)** Light microscopy of differentiated hMADS extruding lipid-like material. **(G)** Toluidine staining of an hMADS cell extruding a lipid vacuole (resin-embedded). **(H)** Quantitative analyses of the amount of material extruded from the cell in SARS-CoV2 infected and uninfected hMADS. Ad: adipocyte. Scale bar: A=0,8 µm, B=120 nm, C=1,2 µm, D= 200 nm, E=40 nm.

**Suppl. Fig. 2 Schematic representation of the prevalence of fat embolism (FE) and lipidic hyaline membranes (HM) in the study population**. Weight status *i*.*e*., OW-OB: overweight and obese subjects with BMI: body mass index (kg/m^2^) ≥25); NW: normo-weight subjects with BMI≤25. Number of patients for each category is reported in parenthesis.

**Suppl. Fig. 3 Fat embolic features in liver of three different COVID-19 subjects with documented lung fat embolism. (A)** Focal, macrovescicular steatosis evidenced by Oil-Red O staining (ORO). **(B)** Several ORO+ lipid droplets into sinusoids (arrows). **(C)** Portal area enlargement of subject shown in B. Note the large lipid droplets into the portal vein lumen (arrow). **(D)** Cluster of lipid rich vacuoles (arrow), like the one found in the mesenteric adipose tissue vein shown in Fig.1K. Scale bar: A=10 mm, B=7 mm, C=8mm, D=13 mm.

