## Supplementary figures and images for "Visceral Fat Inflammation and Fat Embolism are associated with Lung’s Lipidic Hyaline Membranes in COVID-19 patients"

### Supplemental Figue 2

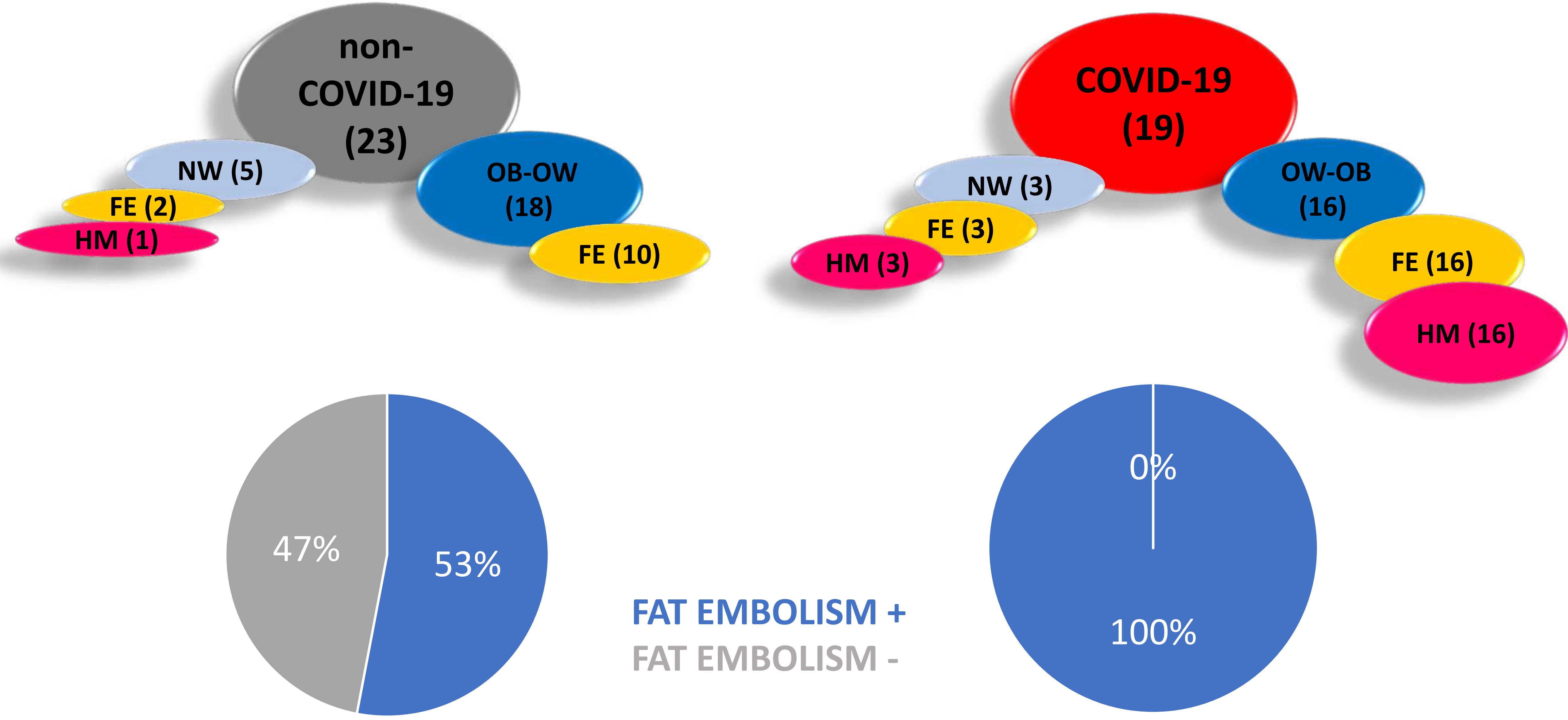

### Supplemental Figue 3

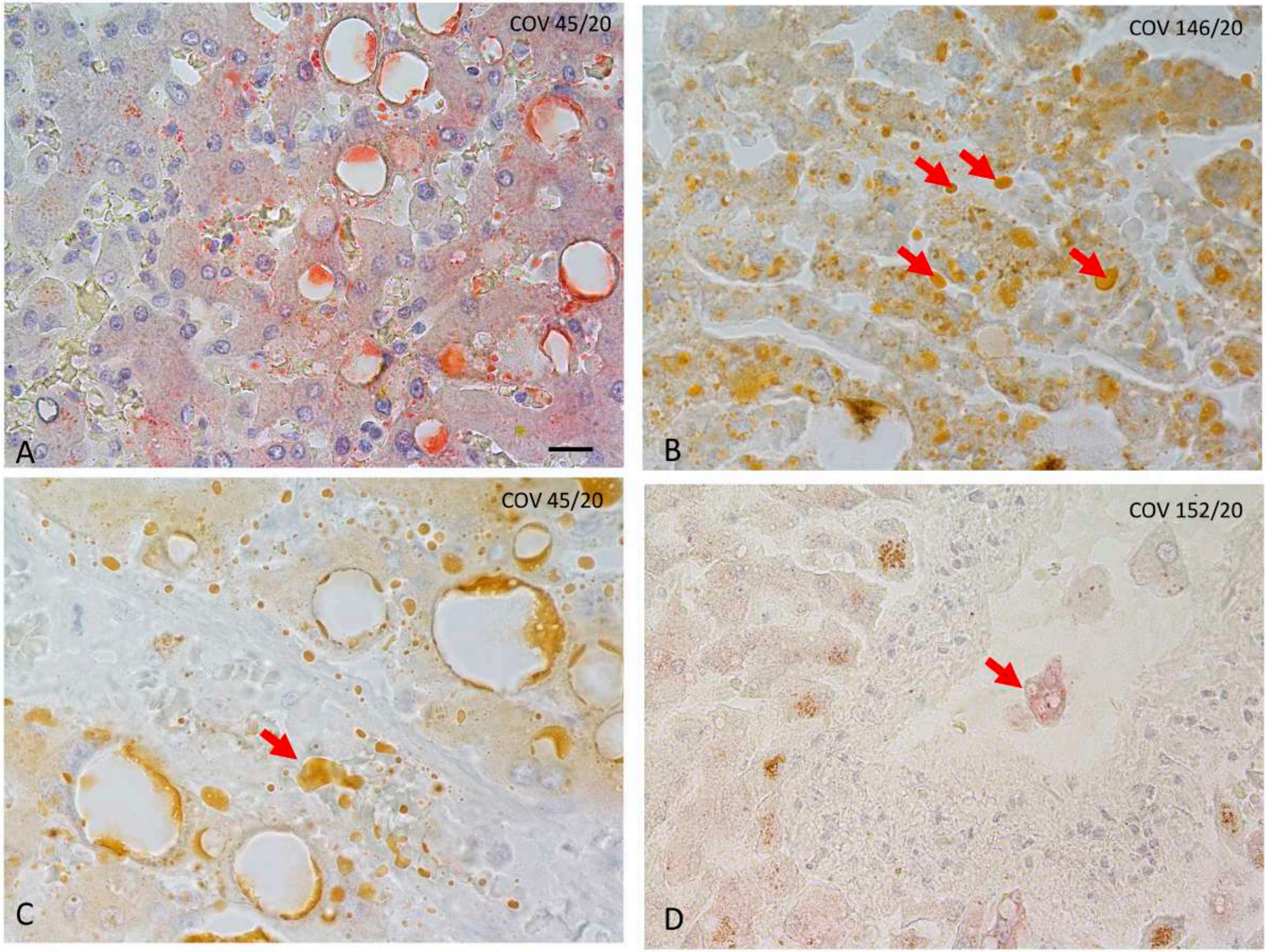

### Supplemental Figure1

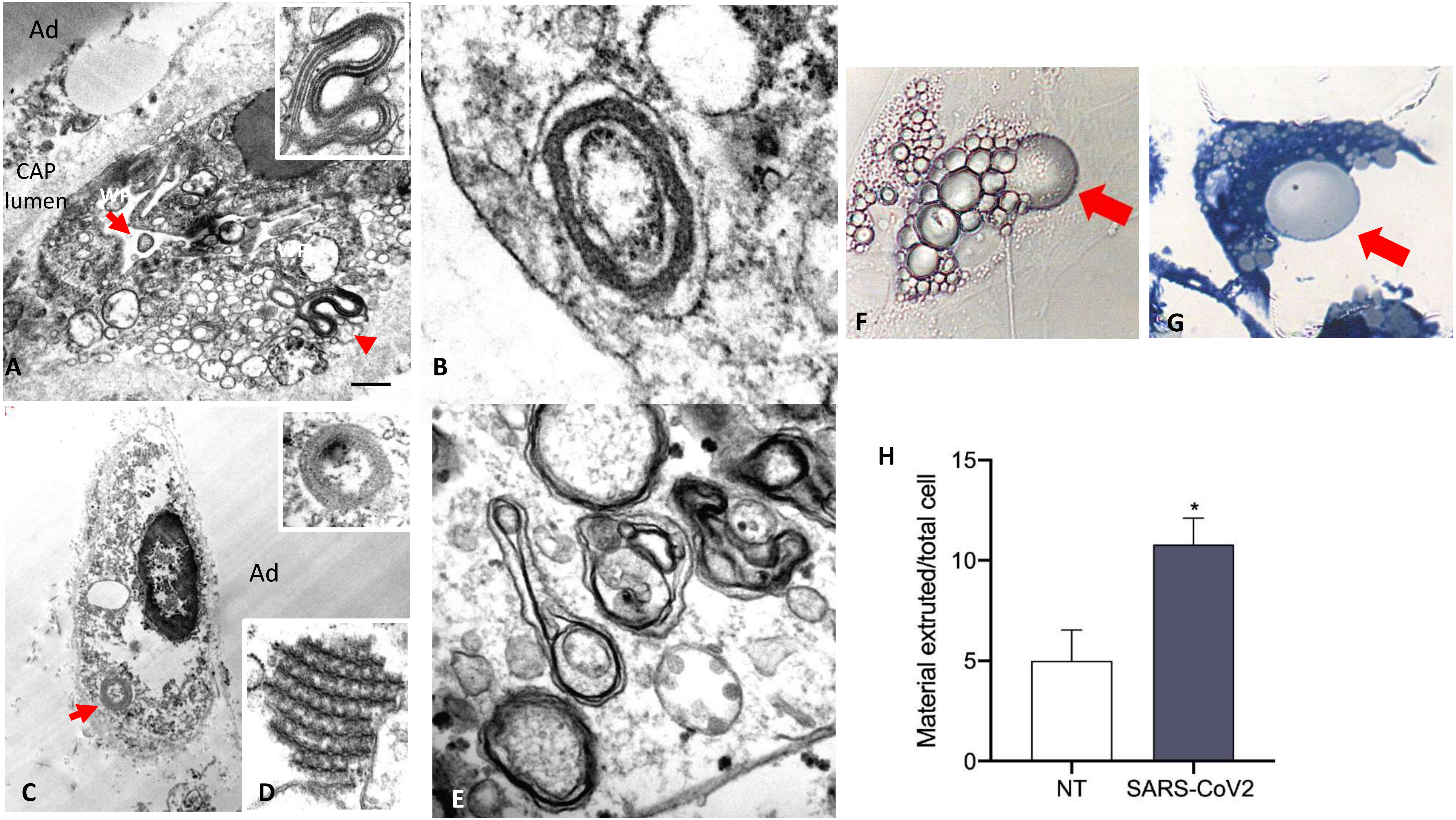
