## Supplemental Tables for "Visceral Fat Inflammation and Fat Embolism are associated with Lung’s Lipidic Hyaline Membranes in COVID-19 patients"

Supplementary Table 1.

| N | ID | SARS-CoV2 by RT-PCR | BMI (kg/m$^2$) | Gender | Age | HPB | T2DM | CVD | PRE-EXISTING RESPIRATORY COMORBIDITIES | CAUSES OF DEATH |
|---|---|---|---|---|---|---|---|---|---|---|
| 1 | 34-18 | - | 28.04 | M | 48 | + | - | + | - | Multiple Organ Dysfunction Syndrome (MODS) |
| 2 | 48-19 | - | 24.2 | M | 74 | - | - | - | + | Respiratory failure |
| 3 | 44/20 | + | 29.3 | F | 62 | + | - | - | + | Cardiac rhythm and/or conduction disease following right myocardial infarction |
| 4 | 45/20 | + | 31.0 | M | 67 | - | + | - | - | Respiratory failure |
| 5 | 50/20 | + | 29.0 | F | 70 | - | - | - | - | Respiratory failure |
| 6 | 52/20 | + | 31.2 | M | 67 | - | - | - | - | Respiratory failure |
| 7 | 56/20 | + | 31.0 | M | 92 | + | - | - | + | Respiratory failure in a setting of pre-existing comorbidities |
| 8 | 67/20 | - | 29.1 | F | 63 | - | - | - | - | Cardiac rhythm and/or conduction disease |
| 9 | 74/20 | - | 27,3 | F | 67 | + | + | - | - | Respiratory failure following pleural empyema |
| 10 | 75/20 | - | 27.5 | M | 58 | + | - | + | - | Cardiac rhythm and/or conduction disease |
| 11 | 78/20 | - | 30.4 | M | 55 | - | - | + | - | Cardiac rhythm and/or conduction disease |
| 12 | 82/20 | - | 20.9 | M | 55 | - | - | - | - | Cardiac rhythm and/or conduction disease |
| 13 | 81/20 | - | 16.0 | M | 27 | - | - | - | - | Lymphoma |
| 14 | 129-20 | - | 29.4 | M | 63 | - | - | - | - | Respiratory depression following acute drug intoxication |
| 15 | 131-20 | + | 30.1 | M | 70 | - | - | - | + | Respiratory failure |

| | | | | | | | | | | |
|---|---|---|---|---|---|---|---|---|---|---|
| 16 | 133-20 | - | 29.3 | M | 72 | - | - | - | - | Cranio-encephalic trauma following accidental fall |
| 17 | 144-20 | + | 28.0 | F | 83 | - | + | - | - | Respiratory failure |
| 18 | 146-20 | + | 31.1 | M | 56 | - | + | - | - | Respiratory failure |
| 19 | 147-20 | + | 27.3 | F | 60 | - | + | - | - | Respiratory failure |
| 20 | 148-20 | + | 24.0 | F | 82 | - | - | - | + | Respiratory failure |
| 21 | 150-20 | - | 28.5 | F | 53 | - | - | - | - | Cardiac rhythm and/or conduction disease |
| 22 | 152-20 | + | 29.4 | M | 51 | - | + | - | - | Respiratory failure and cardio-circulatory arrest. Patient treated with ECMO |
| 23 | 154-20 | - | 31.8 | M | 75 | - | - | + | - | Cardiac rhythm and/or conduction disease |
| 24 | 155-20 | - | 29.4 | F | 44 | - | - | + | + | Cardio-circulatory arrest in pneumonia and metapneumonic pleurisy |
| 25 | 156-20 | - | 31.2 | F | 39 | - | - | - | - | Cardiac rhythm and/or conduction disease |
| 26 | 162-20 | - | 35.5 | M | 43 | - | - | - | - | Cardiac rhythm and/or conduction disease |
| 27 | 00-21 | + | 22.8 | M | 76 | - | - | - | + | Respiratory failure |
| 28 | 0-21 | + | 46.7 | M | 66 | + | - | - | + | Respiratory failure |
| 29 | 8-21 | - | 27.7 | M | 72 | + | + | + | - | Cardiac rhythm and/or conduction disease |
| 30 | 11-21 | - | 30.4 | M | 72 | + | + | + | - | Cardiac rhythm and/or conduction disease |
| 31 | 13-21 | - | 28.3 | M | 55 | - | - | + | - | Cardiac rhythm and/or conduction disease |
| 32 | 17-21 | - | 21.2 | M | 90 | + | + | - | + | Cardio-circulatory arrest in pneumonia and rectal cancer |
| 33 | 28-21 | + | 32.0 | F | 80 | + | - | + | - | Respiratory failure |

| | | | | | | | | | | |
|---|---|---|---|---|---|---|---|---|---|---|
| 34 | 31-21 | + | 36.0 | M | 56 | + | - | - | - | Respiratory failure |
| 35 | 32-21 | + | 23.6 | M | 77 | + | - | + | + | Respiratory failure |
| 36 | 33-21 | - | 45.7 | M | 48 | - | - | + | - | Cardiac rhythm and/or conduction disease |
| 37 | 35-21 | - | 23.4 | M | 79 | - | - | + | - | Carbon monoxide intoxication |
| 38 | 36-21 | - | 25.0 | M | 55 | - | - | + | + | Cardiac rhythm and/or conduction disease |
| 39 | 21/15 | + | 29.4 | F | 71 | - | + | - | - | Respiratory failure |
| 40 | 38-21 | + | 28.4 | F | 53 | - | - | - | - | Respiratory failure |
| 41 | 55/20 | - | 26.3 | F | 58 | + | - | - | - | Cardiac rhythm and/or conduction disease |
| 42 | 03/20 | + | 30.0 | M | 73 | - | - | - | - | Respiratory failure |

Positive symbol indicates presence; negative symbol indicates absence. BMI=Body Mass Index; HBP= High blood pressure; T2DM: Type 2 Diabetes Mellitus; CVD: Cardiovascular disease which includes: cardiopathy, myocardiosclerosis; PRE-EXISTING RESPIRATORY COMORBIDITIES includes: bacterial pneumonia, mesothelioma, amyloidosis; ECMO= Extracorporeal membrane oxygenation.

Supplementary Table 2.

| (n) | Controls (23) | COVID-19 (19) | p |
|---|---|---|---|
| Respiratory conditions*; n (%) | 5 (21.7) | 10 (52.6) | 0.03 |
| Type 2 Diabetes; n (%) | 5 (21.7) | 6 (31.6) | 0.47 |
| Hypertension; n (%) | 9 (39) | 7 (36) | 0.87 |
| Cardiovascular disease; n (%) | 8 (34.7) | 2 (10.5) | 0.06 |

*Pneumonia, dyspnoea, respiratory distress
